# Cell entry inhibitor with sulfonated colloid gold as new potent broad-spectrum virucides

**DOI:** 10.1101/2021.06.11.448146

**Authors:** Chur Chin

## Abstract

We recently developed nontoxic, broad-spectrum virucidal gold nanoparticles, less than 10nm sized, modified with sulfonic acids (mesilate) that mimic heparan sulfates.Camostat, a serine protease inhibitor can introduce gold nanoparticles to the influenza virus via ionic bonds.In this study, we examined the ability of a novel sulfonated colloid gold to inhibit the virus in vivo. Thus, camostat-colloid gold is a promising candidate for the development of antiviral drugs to prevent and treat influenza infection.

## Introduction

Heparan sulfate proteoglycans (HSPGs) are highly sulfated and used by several viruses, for attachment to the cell surface. Becaise of the heavily sulfated chains, HSPGs present a global negative charge that can interact electrostatically with the basic residues of viral surface glycoproteins. HSPG-dependent ACE2 receptor viruses can be grouped as rhino virus, adenovirus, parainfluenza virus, echo virus, Coxsackie virus, corona virus and respiratory syncytial virus (RSV) (1). Gold nanoparticles coated with sulfonic acid inhibit different strains of influenza virus that do not bind HSPGs. The antiviral action is virucidal and irreversible for influenza A (H1N1, H3N2, and H5N1) and B virus strains (2).

We designed antiviral nanoparticles with flexible linkers that mimicking HSPG. They allow effective viral association with a strong binding and multivalent units, generating forces that eventually lead to irreversible viral deformation (3).

## Materials and methods

### Chemicals

Camostat mesylate powder was purchased from Hyperchem (Shanghai, China).Colloid gold (MediGOLD^®^) was purchased from Nutraneering (Irvine, CA, USA) and Tamiflu (oseltamivir phosphate) was purchased from Hoffmann-La Roche Limited (St. Louis, MO, USA).

### Mice and viruses

All research studies involving the use of animals were reviewed and approved by the Institutional Animal Care and Use Committee (IACUC) at the Charles River Laboratory. Six to wight week old female C57BL/6 mice were housed (n=10) and the mouse-adapted influenza strainsA/PR/8/34 (H1N1;PR8) were provided by Charles River Laboratories. The viruses were amplified in 10 day old embryonated hen eggs using standard procedures.

### Infection and treatment of mice

For primary influenza infection, the mice were anesthetized by intraperitoneal injection of Avertin (2,2,2-tribromoethanol; 240mg/kg; Sigma-Aldrich, St. Louis, MO, USA) before intranasal infection with 50 plaque-forming units (PFUs) of PR8 virus in 30mL PBS (Wisent Incorporated). The mice were weighed daily and assessed for clinical signs of the disease. The mice were treated orally twice daily (b.i.d.) during seven days with saline or Tamiflu (oseltamivir phosphate; 5 or 50mg/kg b.i.d.),camostat (30mg/Kg),colloid gold (3mL/Kg) to evaluate morbidity and adaptive immune responses or during 3 day for early cytokine, the immune response, and lung viral titers.

### Organ and cell isolation

Bronchoalveolar lavages (BALs) were used to collect airway cells. Lung were mechanically processed and the cells were isolated through centrifugation and washing steps.

### Lung viral titer determination

Lung viral titers were determined from 10-fold dilutions of clarified supernatants of lung homogenates in viral plaque assays. The number of viral plaques was counted to determine the viral PFUs.

### Real-time PCR analyses

RNA isolated from lung homogenates was reverse transcribed, and real-time PCR was performed. Fold inductions was calculated using the 2-DDC time method and normalized using ribosomal 18S RNA expression. Saline treated and uninfected mice served as the control.

### Cell surface staining and flow cytometric analysis

Fc receptors were blocked with anti-CD16/32 antibody (BD Biosciences, Franklin Lakes, NJ, USA) for 20 min at 4°C. Monocytes/macrophages were identified. Lymphocytes were stained for 30 min at 4°C. After cell staining acquisitions were performed with a FACS Calibur or a FACS Can to flow cytometer, Flow cytometry data were analyzed using the Flow Jo 7.6.4 software (Tree Star, Ashland, OR, USA).

### Cell stimulation and intracellular cytokine staining

Freshly isolated cells (13106/0.2 mL) were stimulated for 6 h with influenza NP366–374 or PA224–233 peptides (1 mM). The cells were subsequently stained for cell surface molecules, fixed, permeabilized, stained for intracellular IFN-g, and analyzed using flow cytometry. Influenza specific CD8+ T cells were identified as IFN-g-producing cells.

### Statistical analyses

The results have been presented as the mean SEM. Data from the four groups were compaired and analyzed using the unpaired Student’s t test with Welch correction. For comparison of four groups, one-way ANOVA followed by Tukey’s post test was performed. Statistical analyses were performed using Graph Pad Prism 5 (La Jolla, CA, USA). A P value, 0.05 was considered statistically significant (*P <.05, **P <.01, and ***P <.001).

## Results

Camostat-colloid gold administration reduces morbidity and decreases influenza replication in mice than Ostelmivir. Camostat was found to be effective in ameliorating mouse influenza by blocking hemagglutinin cleavage (4). In this study, mice were treated with camostat, camostat with colloid gold, oseltamivir and 50 PFU of PR8 virus. The mice that received camostat-colloid gold suffered no weight loss, whereas the mice treated Ostelmivir lost up to 20% of their initial body weight on day 6 post infection (d.p.i.) .Lung consolidation was significantly reduced in Camostat-colloid gold group compared to ostelmivir–treated mice on day 6 d.p.i. There was no difference in the survival rate at d 6 p.i. between ostelmivir-treated and camostat-gold-treated groups. The clinical scores increased over the course of the study in the ostelmivir-treated group, but mostly remained at the base line level in camostat-colloid-gold treated mice (Fig. 1). The symptom score consisted of 1) abnormal gestures of the body 2) pilo-erection 3) abnormal respiration 4) eye discharge and 5) abnormal activity.

**Fig. 1.**
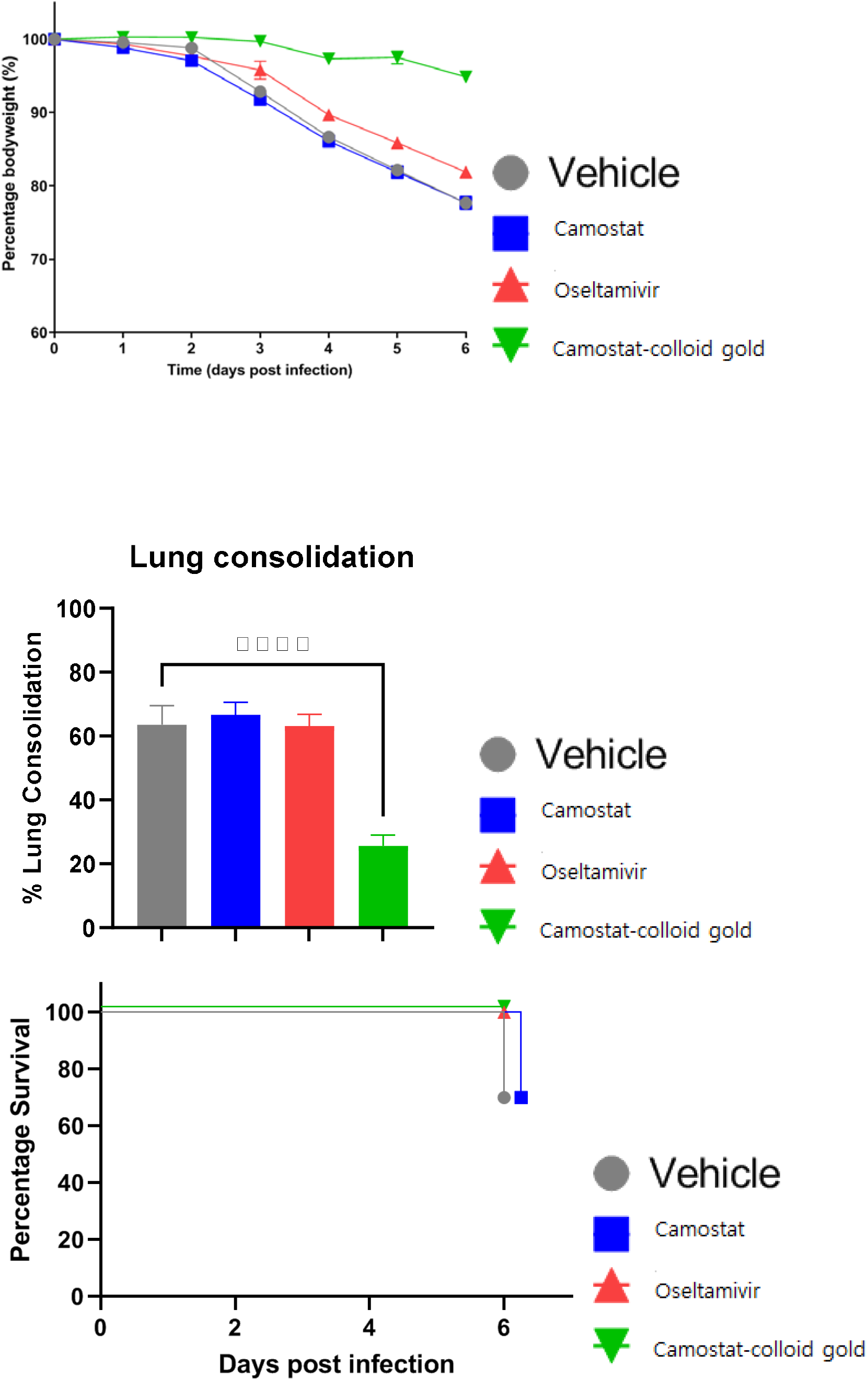
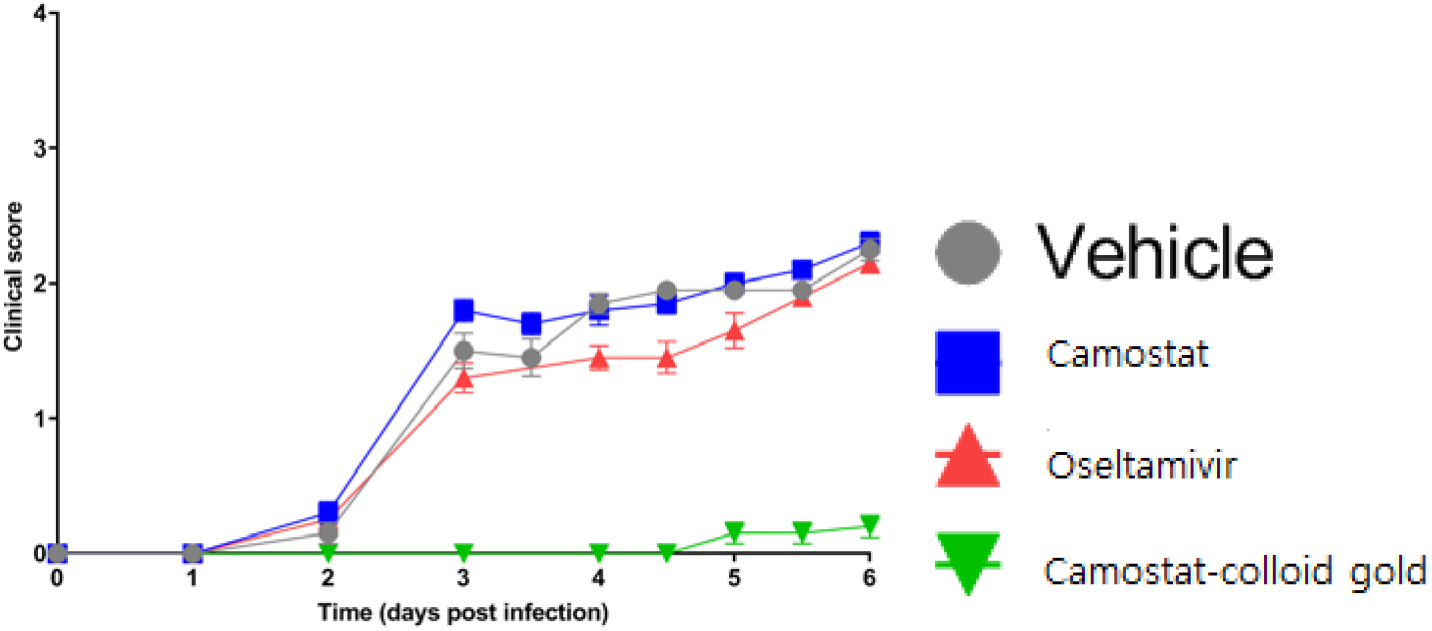
Infection progressed with vehicle treated animals gradually losing body weight and showing increased clinical scores over the course of the study. camostat-colloid Gold reduced body weight loss, clinical score and lung consolidation in the animals.Camostat treatment did not significantly reduce disease progression compared to the vehicle. Oseltamivir treatment resulted in slightly improved clinical scores and survival.

Camostat-colloid gold administration reduced early inflammatory responses in mice during influenza infection compared to to oseltamivir. In general, cytokines levels were higher in BAL than in plasma (5) (Fig.2). Because of the significant effects of camostat-colloid gold administration on cytokine and chemokine expression, we subsequently determined its impact on the recruitment of leukocytes in the lungs as this process is highly dependent on inflammatory chemokines (6).

**FIG 2.**
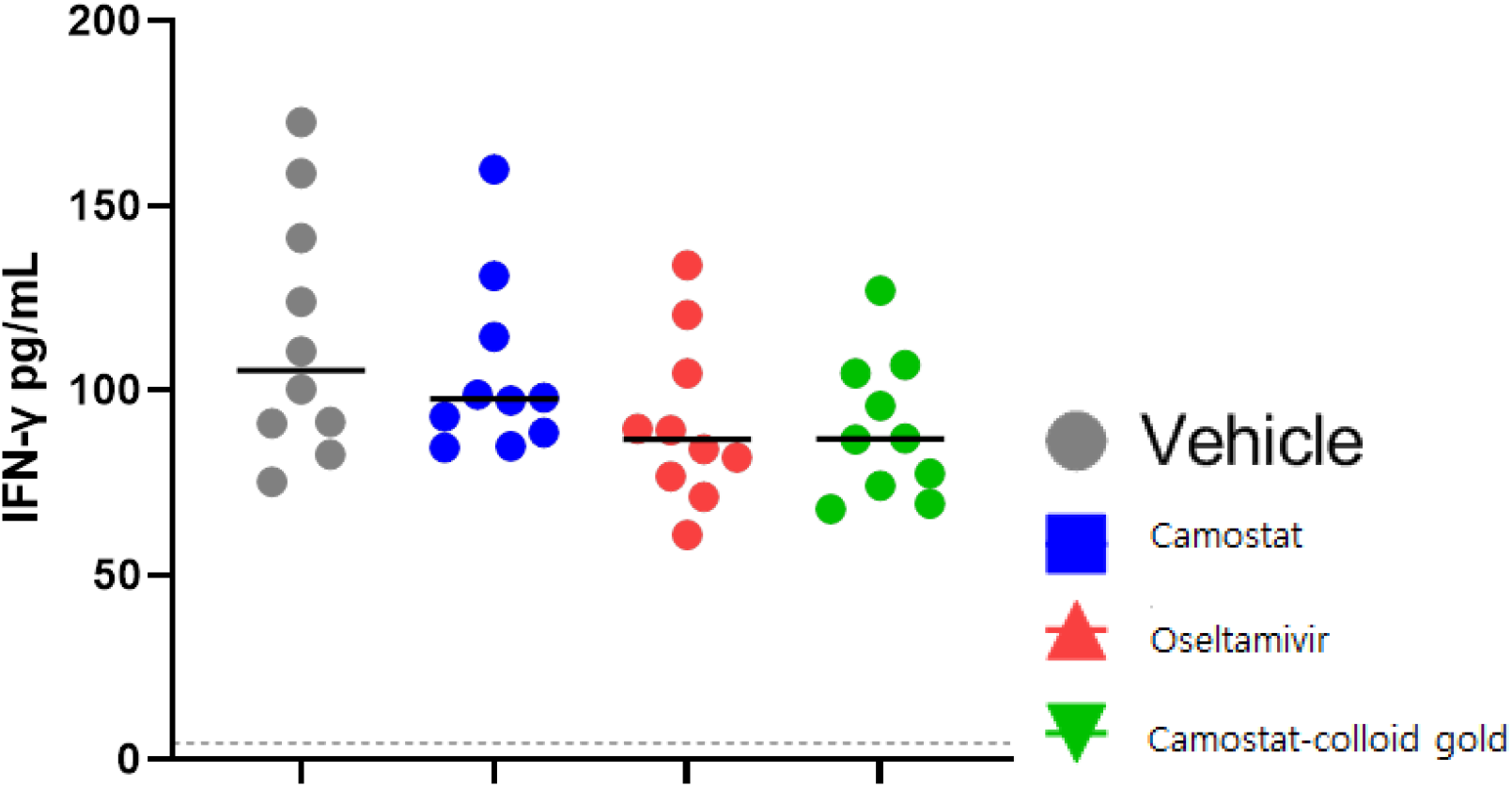
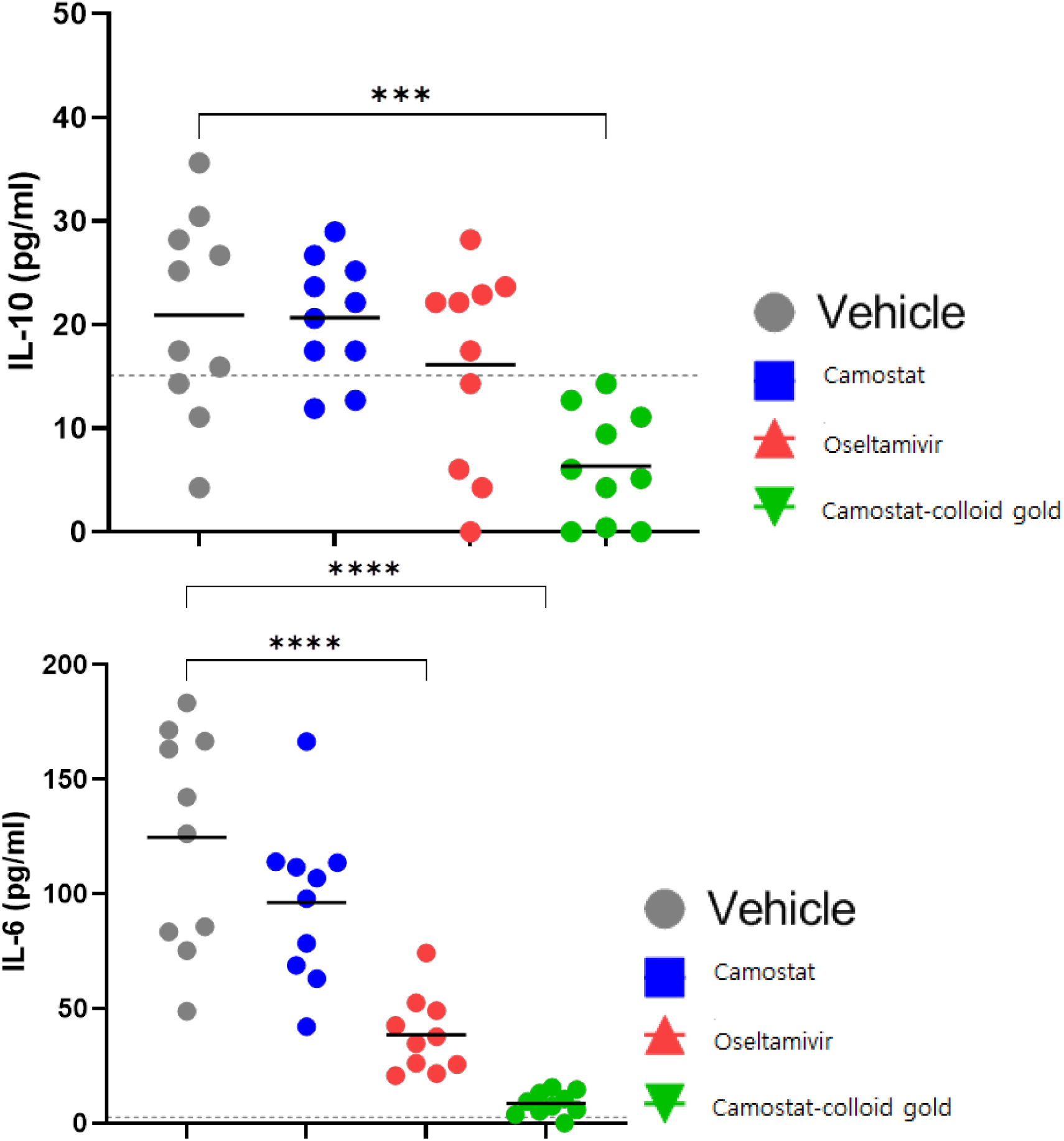
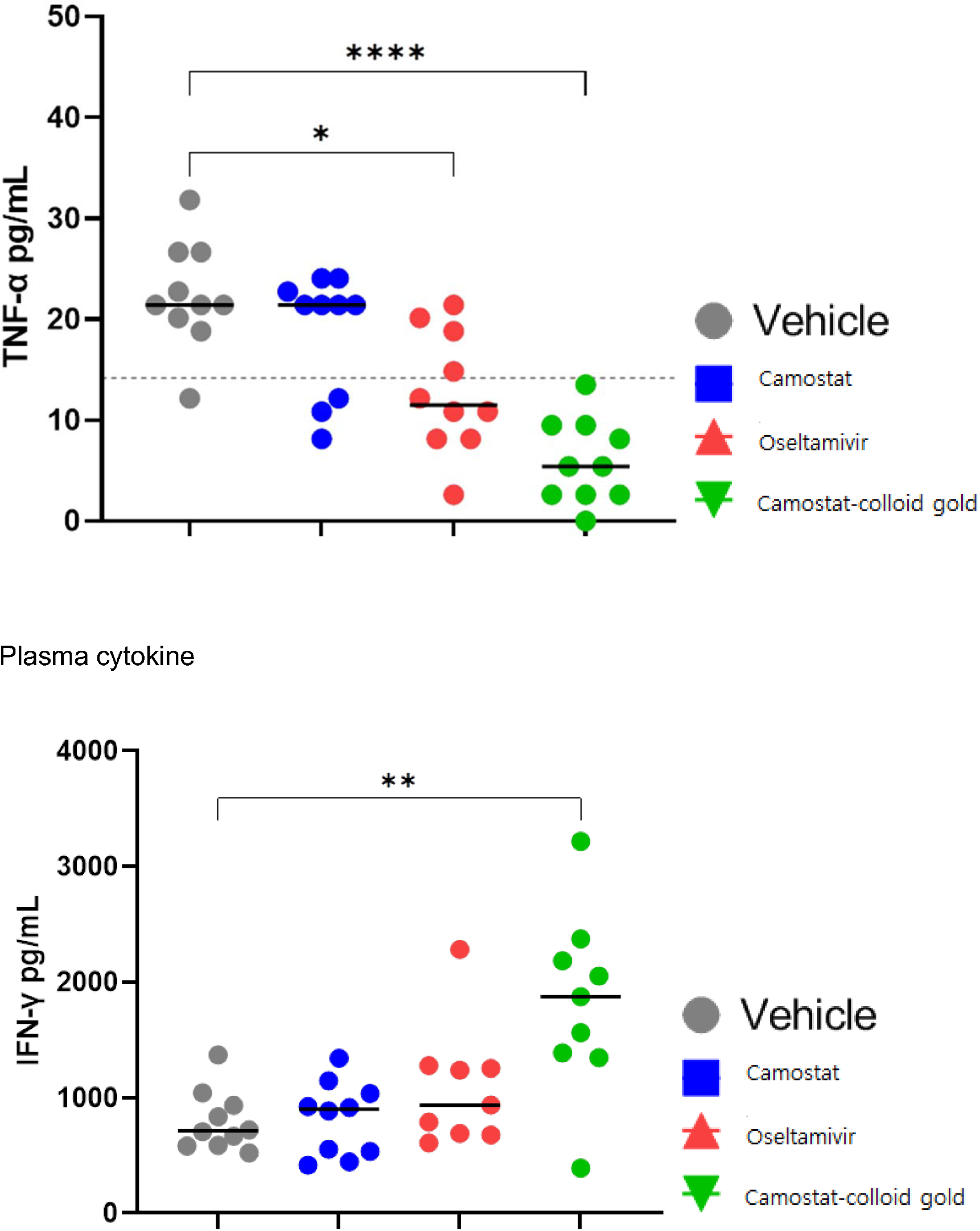
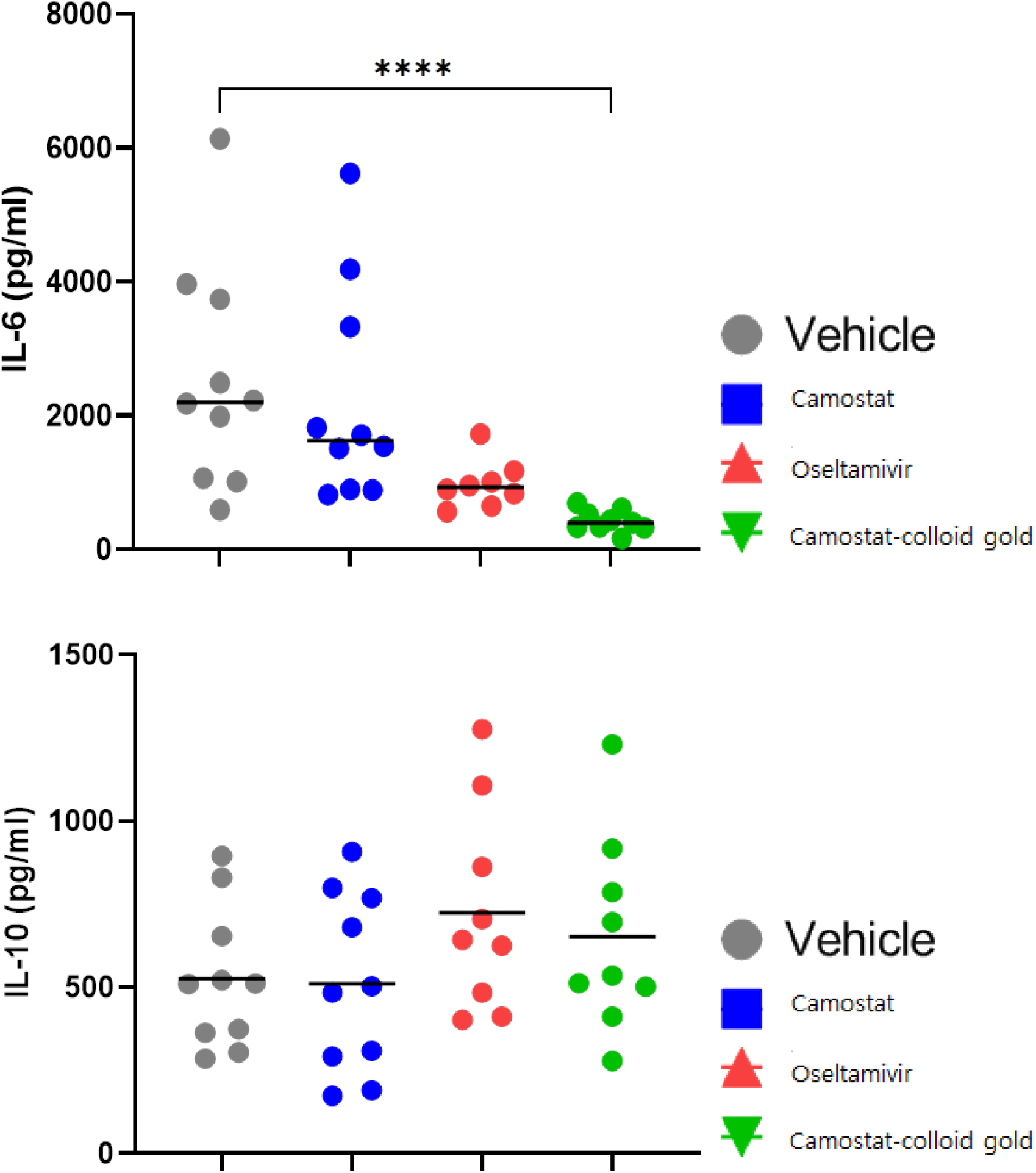
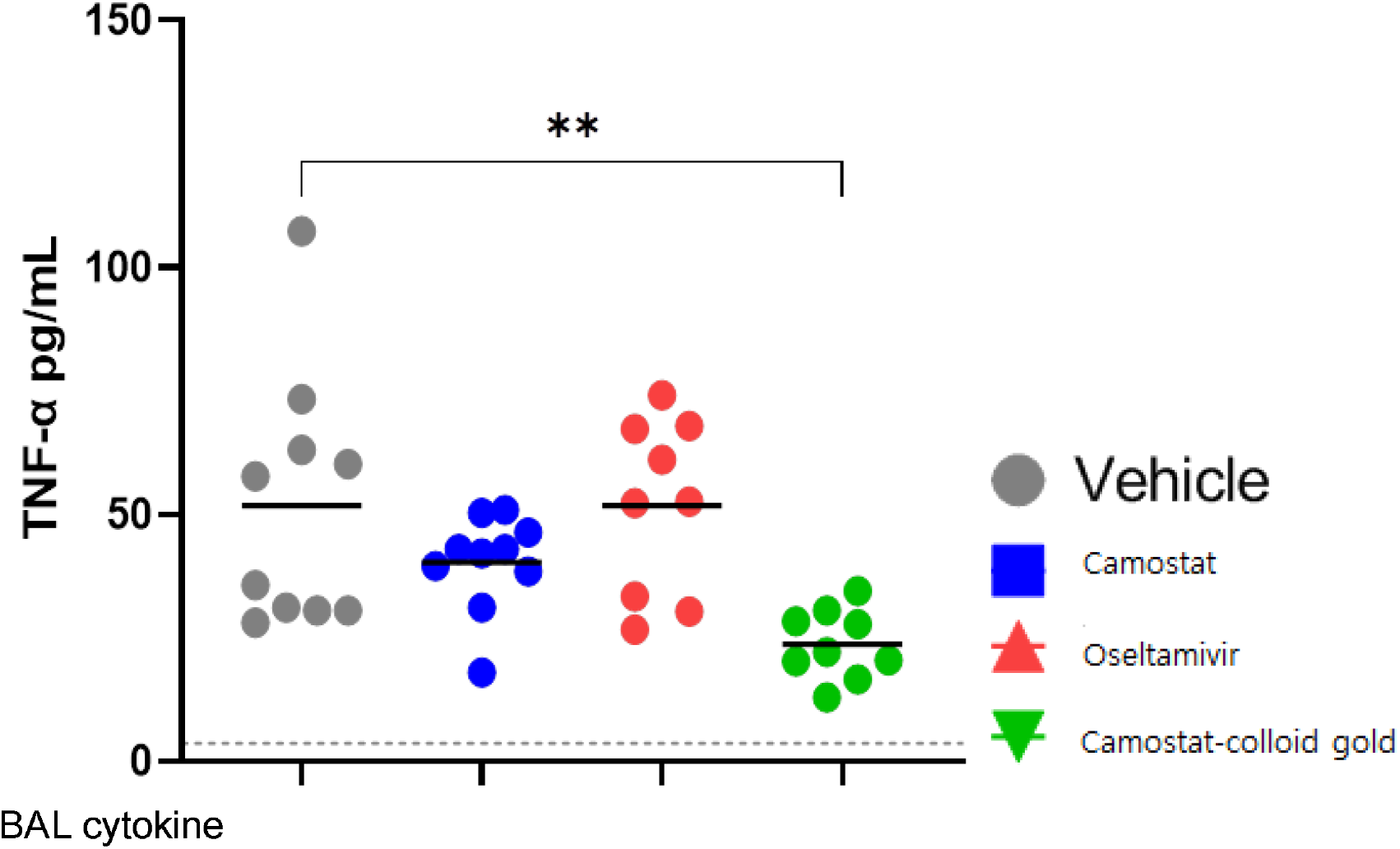
IL-6 (plasma and BAL), TNF-α (plasma and BAL) and IL-10 (plasma) were significantly decreased in the camostat-colloid gold group compared to the vehicle, indicating that the model worked.Oseltamivir induced a statistically significant decrease in IL-6 and TNF-α in plasma (*p*<0.0001 and *p*<0.05, respectively), although levels of TNF-α were predominantly close to or below the functional base line.A decreasing trend was seen in IL-6 levels in BAL for the camostat group compared to the vehicle group, although this did not reach statistical significance. No statistically significant decreases in cytokine levels were seen for the camostat group.

PR8 infection induced a massive recruitment of monocytes/macrophages, neutrophils to the lungs and airways of mice in camostat colloid gold and oseltamivir-treated mice(Fig. 3). The lymphocyte influx into the BALF of influenza infected mice increased (7,8).

**FIG 3.**
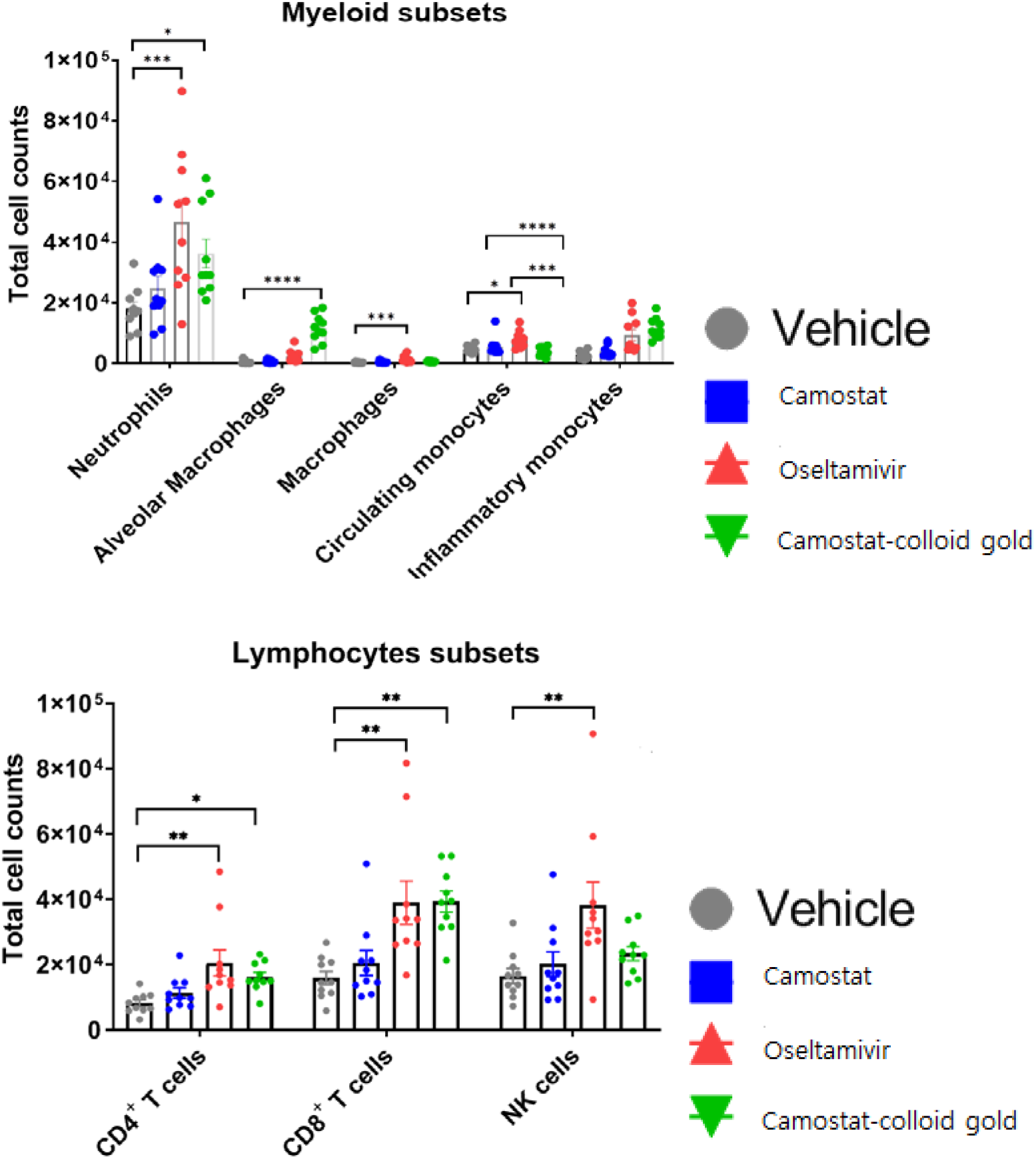
Camostat-colloid gold treatment resulted in a higher count of alveolar macrophages indicating protection against lung injury. The increased infiltration of inflammatory monocytes, CD4 and CD8 cells and T cells in the lungs of camostat colloid gold treated animals by day 6 is indicative of the effect of oseltamivir on slowing disease progression.When compared to vehicle, oseltamivir increased cellular infiltration into the lung for all the cell subsets analyzed except for alveolar macrophages. An increasing trend was seen in the alveolar macrophage count in the BAL in the oseltamivir group compared to that in the vehicle group indicating possible protection against lung injury, although this did not reach statistical significance.

Camostat treatment did not alter immune cell infiltration in the lungs compared to that in the vehicle.

We found that camostat, camostat-colloid gold and osteltamivir caused significantly reduction in lung viral titers.

**FIG 4.**
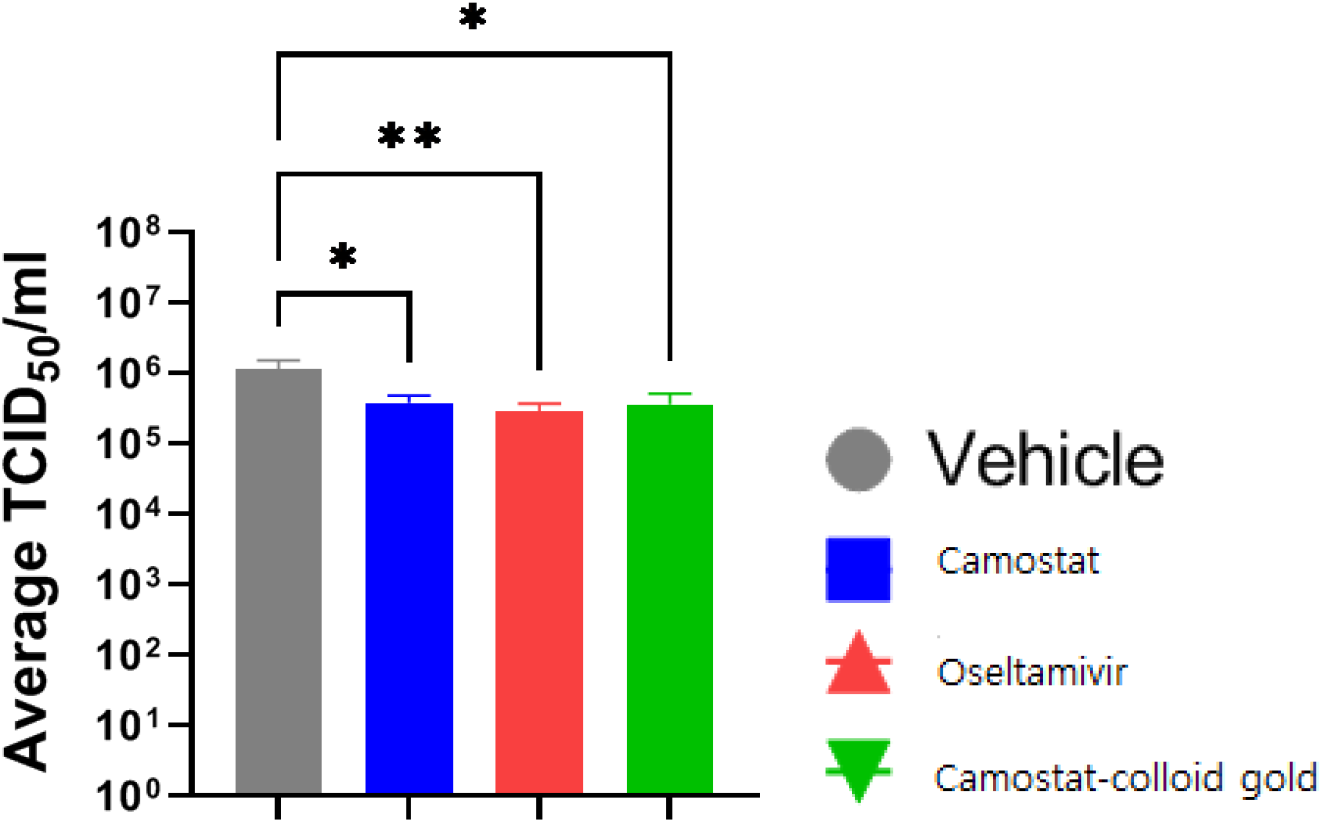
camostat-colloid gold significantly lowered viral load at day 6. Both camostat treatment and oseltamivir significantly reduced the viral load in the lungs at day 6 with camostat-colloid gold.

## Conclusion

Camostat helps ferry gold nanoparticles from the receptor binding sites to the glycoprotein of influenza virus. Gold nanoparticles which are ideal antivirals target conserved viral domains and are virucidal, These properties of gold nanoparticles make them as effective antiviral inhibitor.

## Acknowledgments

The writers would prefer to give their gratitude to the Charles River Laboratory to aid this research.

## Disclosure

The authors declared no conflict of interest for this work.

## Notes

### Competing Interest Statement

The authors have declared no competing interest.

